# Defining Moults in Migratory Birds: A Sequence-based Approach

**DOI:** 10.1101/2021.10.04.463090

**Authors:** Peter Pyle

## Abstract

Two broad nomenclatures have emerged to describe moult strategies in birds, the “life-cycle” system which describes moults relative to present-day breeding and other life-history events and the Humphrey-Parkes (H-P) system which reflects the evolution of moults along ancestral lineages. Using either system, challenges have arisen defining strategies in migratory species with more than one moult per year. When all or part of two moults occur in non-breeding areas they may fail to be recognized as two moults or have been discriminated temporally, whether feathers are replaced in fall, winter, or spring. But in some cases feather replacement can span the non-breeding period, and this has resulted in an inability to identify inserted moults and to compare moult strategies between species. Furthermore, recent analyses on factors influencing the extent of the postjuvenile or preformative moults have either confined this moult to the summer grounds or presumed that it can be suspended and resumed on winter grounds, which has lead to quite divergent results. Evolutionarily, the timing, extent, and location of moults are very plastic whereas the sequence in which feathers are replaced is comparatively fixed. As, such, I propose taking an evolutionary approach to define moults on the basis of feather-replacement sequences as opposed to timing or location of replacement, including strategies in which moults can overlap temporally. I provide examples illustrating the functionality of a sequence-based definition in three migratory North American passerines that can undergo feather replacement twice in non breeding areas, and I demonstrate how this system can effectively apply to moults in many other passerine and non-passerine species. I recommend that authors studying the evolutionary drivers of moult strategies in migratory birds adopt a sequence-based approach or carefully consider replacement strategies both prior to and following autumn migration.

---

Avian moult strategies are complex, as have been terminologies used by ornithologists to describe them. Two broad nomenclatures have emerged to describe bird moults (summarized by Jenni and Winkler 2020:11-14), the “life-cycle” system (Dwight 1900), currently employed by most authors in the Old World, and the Humphrey-Parkes or “H-P” system (Humphrey and Parkes 1959 as modified by Howell et al. 2003), currently employed by most authors in the New World. These two nomenclatures have different bases of definition, the first describing moults relative to present-day breeding and other life-history events and the second defining moults based on how they evolved along ancestral lineages. Among resident and many migratory bird species, especially those that breed in boreal and north-temperate regions, moults are relatively discrete and easy to define based on these terminologies. However, challenges occur defining moults in migratory species, especially those with more than one moult per year, each of which can occur partially or entirely in non-breeding areas.

Among those using the life-cycle terminology, differences of opinion exist about whether the postjuvenile and/or postbreeding moults can begin on summer grounds and resume on winter grounds (e.g., Cramp 1988, for the most part) or are restricted to summering grounds, with feather replacement on wintering grounds being assigned to prebreeding moults (e.g., Jenni and Winkler 2020, for some but not all species). These differences have resulted in confusion in describing life-cycle moults in such species as Barred Warbler *Curruca nisoria* or those of the Red-backed Shrike *Lanius collurio* complex including Isabelline *L. isabellinus* and Brown *L. cristatus* shrikes, species which show evidence of moulting feathers twice on wintering grounds (Stresemann and Stresemann 1971, Cramp 1988, Hall and Tullberg 2004, Kiat and Perlman 2016, Jenni and Winkler 2020, Kiat and Izhaki 2020). Considering such moults on winter grounds as a single prebreeding moult compromises our ability to identify inserted moults and to discriminate and understand moulting strategies. Furthermore, recent analyses on factors influencing the extent of the postjuvenile and preaternate moults in passerines have either confined this moult to the summer grounds (Guallar and Figuerola 2016, Delhey et al. 2020, Pérez-Granados et al. 2020, Kiat et al. 2021) or presumed that it can resume on winter grounds (Guallar et al. 2021), which has lead to quite divergent interpretations of results.

The H-P system is often portrayed as attempting to define evolutionary homologies (e.g., Jenni and Winkler 2020) whereas it is better considered as reflecting the end product of moult evolution itself. Although these two characterizations may appear to differ only subtly, the former interpretation would presume guesswork whereas the latter solidifies the definition of moults, despite the fact that their evolutionary histories are only starting to be elucidated with phylogenetic analyses (e.g., Kiat et al. 2019, Terrill et al. 2020, Guallar et al. 2021). The definitive prebasic moult is best regarded as having evolved from ecdysis during an annual (or cyclic) restorative process in reptiles (Howell and Pyle 2015, Kiat et al. 2020), and it may be possible that the preformative moult has also evolved from an extra inserted ecdysis event in younger reptiles as their body size grows. These moults are present in the great majority of birds and can be considered ancestral under H-P nomenclature. Within this framework, additional prealternate moults and occasionally third presupplemental moults have evolved and become inserted within the basic cycle (Humphrey and Parkes 1959). Importantly, prealternate and presupplemental moults have emerged at different times along ancestral bird lineages, and are thus not considered homologous throughout all birds (Howell et al. 2003), just within each linage from the time the moult first evolved. Such an evolutionary approach can be applied to birds globally rather than favouring boreal-breeding species that form the basis of other terminologies.

Using this approach, moults that commence on or near breeding grounds and complete at stopover moulting grounds (cf. Pyle et al. 2009, 2018a) or on winter grounds are recognized evolutionarily as single moults. Substantial variation in moult timing, location, and extents of moults before and after suspension can occur among closely related species, within species such as Brown Shrike (Cramp 1988), Common Whitethroat *Currica communis* (Jenni and Winkler 2020), and Warbling Vireo *Vireo gilvus* (Voelker and Rohwer 1998), and even within the same individual interannually (Pyle et al. 2009). It is thus clear from an evolutionary perspective that the timing and locations of moults relative to migration are relatively recent adaptive responses to life-history traits and environmental factors (Pyle et al. 2018a, Kiat et al. 2019, Delhey et al. 2020, Terrill et al. 2020, Guallar et al. 2021), rather than their being separate moults that have evolved independently at different locations. It follows that the various terminologies used to describe minor or perceived differences in spatial and temporal replacement patterns in some (but not necessarily all) individuals of a species, including “split moults,” “interrupted moults,” “seasonally divided moults,” “interlaced moults,” and in some cases “arrested moults,” may represent variations upon a single evolutionarily derived strategy, broadly termed “suspended moults” by Jenni and Winkler (2020) and those now using H-P nomenclature (e.g., Pyle 2008, Tonra and Reudink 2018).

## IDENTFYING AND INTERPRETING MOULTS

Under either terminology there remains uncertainty in discriminating preformative/postjuvenile or prebasic/postbreeding moults from prealternate/prebreeding moults when feathers are replaced more than once away from breeding grounds. For example, Pyle (1997) considered protracted overwinter flight-feather replacement in both first-year and adult Yellow-bellied Flycatcher *Empidonax flaviventris* and several other species to include prealternate moults whereas a similar replacement strategy in Red-eyed Vireo *Vireo olivaceus* and many waterbirds (Pyle 2008) was considered only to include the prebasic moult. Similarly, Jenni and Winkler (2020) consider moults of some adult swallows, pipits and some populations of Common Whitethroats to be a single suspended postbreeding moult whereas similar moults in, for example, Barred Warbler, other populations of Common Whitethroats, and European Pied Flycatcher *Ficedula hypoleuca* were considered separate postbreeding and prebreeding moults. In some cases, partial post breeding moults on summer grounds followed by complete prebreeding moults on winter grounds have been reported (e.g., for *Acrocephalus* and *Phylloscopus* warblers, Garden Warbler *Sylvia borin*, and Spotted Flycatcher *Muscicapa striata*), but documentation of moults on winter grounds is widely considered sparse, and evidence that these species are replacing feathers twice rather than suspending moults (in some but not all individuals) is thus far inconclusive (Jenni and Winkler 2020). Indeed, further consideration of wear patterns in the feathers of Arctic Warbler *Phylloscopus borealis* suggested the possibility that only one complete and suspended moult occurs per year (Snyder 2008), rather than a partial moult followed by a complete moult as previously reported (Cramp 1992).

## A SEQUENCE-BASED APPROACH

Evolutionarily, the timing, extent, and location of moults are very plastic whereas the sequence in which feathers are replaced is comparatively fixed (Pyle 2013). During prebasic moults of remiges, the majority of birds follow the “basic sequence” (Ginn and Melville 1983), replacing primaries distally from a node at the innermost primary (P1; see Table 1 for feather numbering) and secondaries bilaterally from a node at the second tertial (T2 in non-passerines or S8 in passerines), proximally from the outermost feather (S1), and in larger diastataxic species, proximally as well from S5 (Pyle 1997, 2005, 2008, 2013; Rohwer 2008; Jenni and Winkler 2020). These sequences are well documented in resident bird species and are maintained in those migratory species in which sequence has been studied. When exceptions to these sequences occur, they are usually found among all individuals and species throughout a lineage once evolved (Pyle 2013), for example among avian orders Gruiformes, Falconiformes, and Psittaformes, families such as Diomedeidae, and passerine species such as Spotted Flycatcher (Pyle 2008, 2013; Rohwer and Rohwer 2018; Jenni and Winkler 2020). Rare exceptions to fixed remigial sequences among passerine genera such as *Lanius, Locustella, Rhipudura*, and *Muscicapa* (Cramp 1988, Junda et al. 2012, Kiat 2017, Jenni and Winkler 2020) are perhaps best considered recent or current evolutionary divergences within these taxonomic lineages.

**Table 1.** Examples of sequence-based definitions following the preformative and first prealternate moults in first-year male Summer Tanagers *Piranga rubra* and Indigo Buntings *Passerina cyanea*. Links are to images catalogued by the Cornell Lab of Ornithology’s Macaulay Library. Feather-tract definitions, feather numbering, and abbreviations for marginal coverts (MaC), median coverts (MeC), greater coverts (GC), carpal covert (CC), alula (A), secondaries (S), and primaries (P) follow those of Figure 9 in Jenni and Winkler (2020:10); rectrices (RR) are numbered proximally on each side of tail. Designations are proposed for illustration of a sequence-based approach and, in some cases may be ambiguous, approximate, or not listed if feathers are not entirely diagnosible.

| Macaulay # | Formative Summer | Formative Fall/Winter | Formative Spring | First Alternate Spring |
| --- | --- | --- | --- | --- |
| A. Summer Tanager |  |  |  |  |
| ML62817821<sup>1</sup> | MaC, GC8-9, A1-2 | MeC, GC1-7, 10 |  |  |
| ML157088011 | MeC1-3, A1-2 | MeC4-8, GC1-8 | S8-9, RR | GC9-10 |
| ML233637131 | MaC, MeC, GC2-7, A1-2 |  | GC1, S8-9, RR | MaC, MeC6, GC8-10 |
| ML55755651 | some MaC, MeC1-4, A1 | GC1-5, CC, A2, R5-6 | S8-9<sup>2</sup>, R3-6 | some MaC, MeC5-8, GC6-10, R1-2 |
| ML153839791 | Mec8, A1 |  | S8-9<sup>3</sup>, P1-2<sup>3</sup>, RR | MaC, MeC, GC, RR |
| B. Indigo Bunting |  |  |  |  |
| ML51314551 | GC6-7, S8-9 | GC1-5, S5-6, P5-9, RR | A2-3 | Mac, MeC, GC7-10 |
| ML236845401 | MeC8, GC1, 3; S7-9 | A2-3, P6-9, RR |  | MaC; MeC; GC2, 4-10 |
| ML94449631 | GC4-5, S8 | P5-9, S9, RR | some MaC, GC1-3, CC, S7 | MaC, MeC, GC6-10 |
| ML59025811 | some MaC; GC1, 9-10; S9 | GC7-8, S7, P4-9, R2-6 | S4-6 | Mac, MeC, GC2-6, S8, R1 |
| ML157869921 | GC1-3, 10; MeC5, S7,9, R3-6 | S4-6, A2-3, P5-9 |  | MaC, MeC, GC4-9, S8, R1-2 |

<sup>1</sup>Image taken in December; remaining images taken during 14 April-19 May.
<sup>2</sup>S8 may be alternate
<sup>3</sup>Adventitious or anomalous preformative replacement pattern for this species

Remigial replacement during preformative moults is likewise fixed, following the same sequences described above for prebasic moults, despite varying nodal positions among primaries and a proximal wave from S1 appearing to be suppressed in many passerines exhibiting eccentric moults (Pyle 1997, Gargallo 2013, Jenni and Winkler 2020). In these and other species exhibiting partial but not eccentric preformative moults, distal replacement from a node at the second tertial appears to be prioritized among secondaries, with for example S6 through S4 being replaced distally and at decreasing rates of frequency (Pyle 1997, Gargallo 2013); replacement extent during eccentric moults has been correlated with hatching date within the breeding season (Elrod et al. 2011).

Among upperwing secondary coverts, feathers are generally replaced in a distal direction among each tract (marginal, median, and greater coverts) during preformative moults, as implied by most resultant moult phenologies (Guallar and Jovani 2020a, 2020b), whereby most or all formative feathers are located proximal to most or all juvenile feathers, defining moult limits following partial replacement (Pyle 1997, Jenni and Winkler 2020). Although exceptions and some variation in covert-replacement phenologies can occur between and occasionally within species (Pyle 1997, Guallar et al. 2014, Guallar and Jovani 2020b, Jenni and Winkler 2020), in most cases a prevailing order to feather replacement appears to be maintained. Prealternate moults may involve different replacement mechanisms, as feather-replacement sequences can vary to a greater extent than those of prebasic and preformative moults, e.g., among the secondaries of gulls (Pyle et al. 2018b) and the resultant wing-covert phenologies of passerines (Guallar and Jovani 2020a, 2020b; Jenni and Winkler 2020). In most cases, however, replacement of coverts is less extensive during prealternate than during preformative moults and the sequences of each can be identified (see examples in Figures 1 and 2 and Table 1, and discussion of exceptions to this pattern below). Among prebasic and preformative moults within species, by contrast, sequence is relatively fixed among broader evolutionary lineages, and may be controlled by neurological as opposed to hormonal mechanisms within restorative processes that include moult (Pyle 2013).

**Figure 1.**
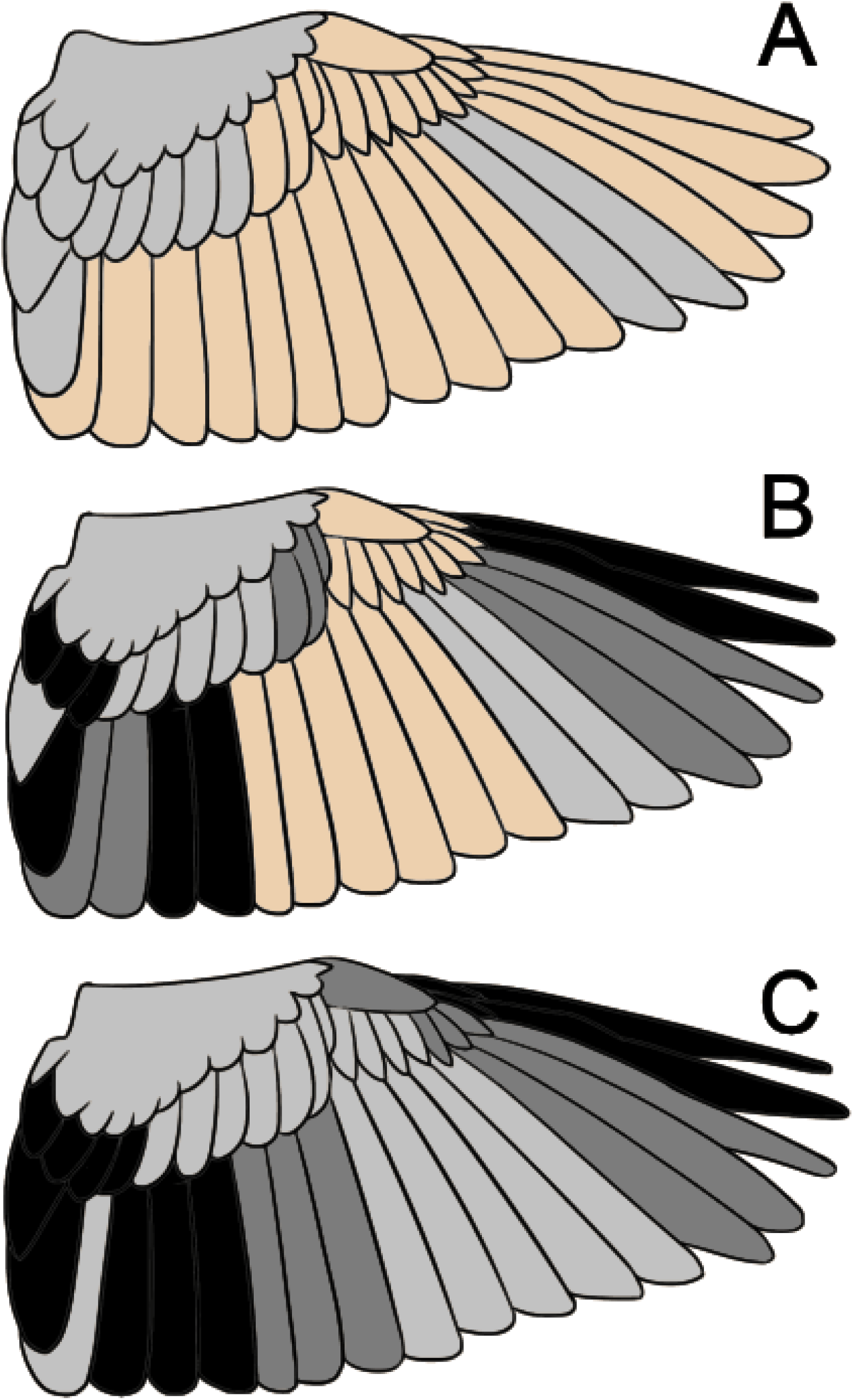
Progression of preformative, prebasic, and prealternate moults away from breeding grounds in Western Kingbird (*Tyrannus verticalis*); see Table 1 for feather numbering. (**A**) Formative feathers (pale gray) replaced in late summer on moulting grounds include inner greater coverts, S8-9, and P6-7, before suspending eccentric moult for migration to winter grounds; remaining feathers are juvenile (tan). (**B**) Similar to (A) but preformative moult had continued on winter grounds with outer greater coverts, S6-7, and P6-8 (dark gray) in fall and winter, and with S5-4 and P9-10 in spring (black), overlapping first prealternate moult of inner three greater coverts and S8 in timing (also black). (**C**) Adult in summer following definitive prebasic and prealternate moults: basic secondary coverts, S7-9 and P1-5 along with corresponding primary coverts were replaced in late summer on moulting grounds, followed by suspension for migration, replacement of S1-3 and P6-8 in fall and winter, and replacement of S4-6 and P9-10 and corresponding primary coverts in spring (black) overlapping prealternate moult of inner greater coverts and S8-9 in timing (also black). Wings based on specimens at the California Academy of Sciences (CAS): (**A**) CAS63960, Guerrero, Mexico, 7 November 1950; (**B**) CAS 88236, California, 28 May 1965; (**C**) CAS46917, California, 12 June 1898. See also Figure 172 in Pyle (1997) for more information and see <u>PSM17928</u> at Slater Museum of Natural History (2021) for an example with moult patterns similar to (**C**).

**Figure 2.**
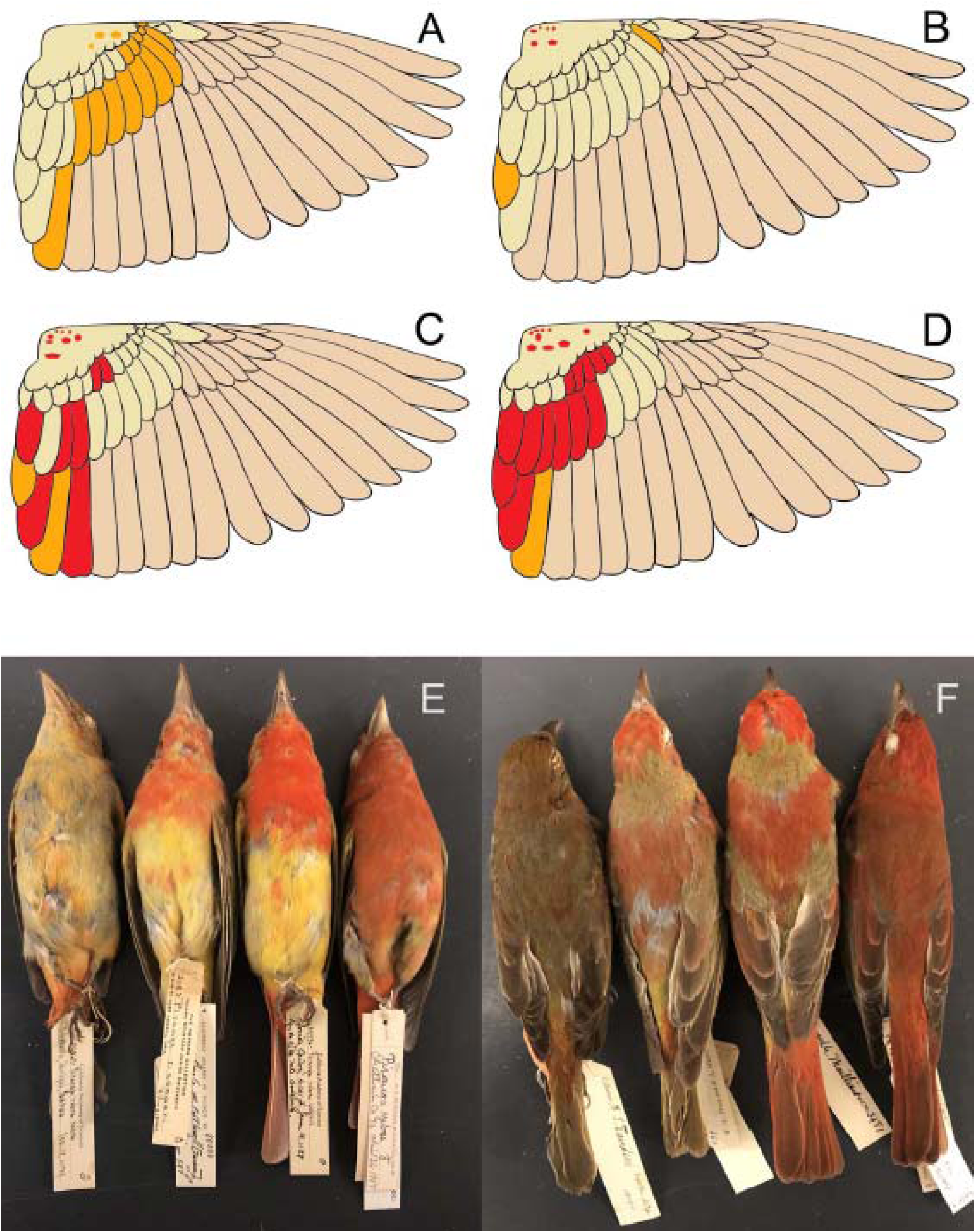
Progression of peformative and first-prealternate moults in first-year male Summer Tanagers (*Piranga rubra*) as based on specimens at the California Academy of Sciences (CAS). Primaries, primary coverts, and most secondaries remain as juvenile feathers (tan) until the second prebasic moult. (**A**) Formative feathers were replaced on or near summer grounds (yellow) followed by suspension for migration and continued replacement of formative feathers in late fall or early winter on winter grounds (CAS32863, Panama, 10 December 1929). (**B**) Most formative secondary coverts and S8 were replaced on or near summer grounds (yellow), followed by formative carpal covert and S9 replaced in fall or winter (orange) and first prealternate body feathers (see below) and marginal coverts (red), replaced in spring (CAS80003, Arizona, 19 June 1893). (**C**) Similar to (**B**) but both S7 and S9 were replaced in spring (red) and some median and greater coverts and S8 are first alternate, also red and replaced in spring; the S8 was presumed also replaced during the preformative moult (CAS29736, Arizona, 16 June 1927). (**D**) Similar to (**C**) but S6 was replaced at the end of the preformative moult in spring, overlapping the first prealternate moult that included a greater number of wing coverts and S8-9 (CAS53100, Georgia, 26 April 1907). (**E** and **F**) The above four specimens (left-to-right), ventral and dorsal aspects, respectively, showing formative (yellow, orange, and red) and first alternate (red) body feathers with feather colour generally reflecting replacement locations and timing (cf. Figure 22 in Howell 2010:17). The left two individuals had retained all juvenile rectrices whereas the right two birds had formative rectrices that were replaced in late winter or spring. Undertail coverts and uppertail coverts were often orange, indicating retention of juvenile feathers in this tract over fall migration. See Table 1 for feather numbering and more examples of first-cycle males.

As, such, I propose defining moults that take place partially or entirely on non-breeding grounds on the basis of evolutionarily fixed feather-replacement sequences as opposed to highly plastic timing or location of replacement. Sequential feather replacment can be suspended for migrations and protracted throughout nonbreeding periods, leading to strategies in which partial (prealternate) body-feather and wing-covert moults, commencing a new sequence, can overlap the completion of protracted flight-feather and wing-covert moults.

For example, in the well-studied Western Kingbird *Tyrannus verticalis*, moult of primaries in both first-year birds and adults can variably commence on breeding grounds, occur partially or entirely at stopover locations, and/or suspend to complete as late as spring on winter grounds, with a second partial replacement of feathers occurring on the winter grounds in spring (Pyle 1997, Rohwer 2008, Barry et al. 2009). Under the sequence-based definition proposed here, the entirety of remigial replacement, despite where it occurs, is considered part of the prebasic or preformative moult, with an overlapping prealternate moult in spring that includes a renewed sequence of replacement among body feathers, inner greater coverts, and tertials (Figure 1). Eccentric preformative moult of primaries usually completes starting from a predetermined node (Gargallo 2013) and thus should be considered a single moulting episode as opposed to two separate episodes, as previously interpreted by Pyle (1997).

Due to gradual temporal shifts in plumage colouration during protracted moults, first-year male Summer Tanagers *Piranga rubra* present a good example for defining first-cycle moults based upon sequence among upperwing coverts and tertials. Adult males in definitive basic and definitive alternate plumages exhibit uniformly red feathers whereas in first-cycle males, feather colour appears to track timing of replacement, juvenile feathers being dull yellow, formative feathers replaced in late summer and early fall being brighter yellow, and feathers replaced on winter grounds varying from orange-yellow in late fall to red in spring (Figure 2). Although feather colour should not be used to define moults (Howell et al. 2003, Howell 2010), the progression of the partial preformative and prealternate moults in first-year male Summer Tanagers can be assumed according to the extent of redness in feather colouration (cf. Figure 22 in Howell 2010:17), and indicates that this moult can occur throughout the non-breeding period and overlap a prealternate moult in spring (Figure 2, Table 1A). Moult of brown and blue wing feathers can similarly be traced with first-year male Indigo Buntings *Passerina cyanea* (Table 1B). In both the kingbird and tanager examples, first prealternate moults can be elucidated and, as expected, are found to be similar in extent to that of adults in passerines (Pyle 1997).

## CLEARING UP PREVIOUS INTERPRETATIONS

Taking a sequence-based definition of moults has the potential to clear up previously confused terminologies in species among the Red-backed Shrike complex (Pyle et al. 2015) and Yellow-bellied Flycatcher (Carnes et al. 2021), allowing for direct comparison of moults among these genera. Among European species treated by Jenni and Winkler (2020), moulting episodes can also be clarified. Barred Warblers, for example, can be considered to undergo a complete prebasic (postbreeding) moult and an incomplete eccentric preformative (postjuvenile) moult, each of which concludes with secondaries on winter grounds; this interpretation may help confirm suspicions that eccentric moults are confined to the first cycle. The “additional prebreeding moult” reported to include wing coverts, tertials, and rectrices, would be considered a new moulting sequence involving the prealternate (prebreeding) moult which, in some individuals, overlaps the conclusion of the suspended moults. Similarly, for first-year Western Yellow Wagtails *Motacilla flava* the “first phase” of the prebreeding moult would be considered part of a suspended preformative moult while the “second phase” of the prebreeding moult would be considered the prealternate moult in both first-year and adult birds, as defined by a renewed sequence in feather replacement. If a third replacement of body feathers occurs in this species, as sometimes reported but may not be fully documented at the individual level (Cramp 1992), a presupplmental moult would be identified under H-P terminology, and its placement would depend on when it evolved along ancestral lineages. In Figure 564 of Jenni and Winkler (2020), greater coverts 4-5 could be formative and greater coverts 6-10 juvenile, rather than all being a single generation (prebreeding). All populations of Common Whitethroats would undergo complete prebasic and partial (occasionally eccentric) preformative moults which, as in *Lanius* shrikes (Pyle et al. 2015), can vary as to geographic location and extent prior to suspension for southbound migration. Potential extra inserted prealternate moults on winter grounds (e.g., in Figures 313-320 of Jenni and Winkler 2020) can also be detected, and variation in moult strategies in Common Whitethroats can be compared in an evolutionary sense. Inserted prealternate moults can similarly be identified in other species with moult categories 3-5 of Jenni and Winkler (2020:34-38, 72-74) including, for example, the *Curruca* warblers illustrated in Figure 57, which appear to have undergone eccentric preformative moults and have first alternate tertials (S8-S9) and inner greater coverts.

In many species, a sequence-based definition results in the prealternate moult being defined as simply including those feathers moulted for a second time within the first moult cycle; i.e., replacing either formative or basic with alternate feathers, but this does not form the basis of definition. In some species the first prealternate moult can be more extensive than the preformative moult, for example in Willow Warbler *Phylloscopus trochilus*, Indigo Bunting, American Yellow Warbler *Setophaga petechia*, and Bobolink *Dolichonyx oryzivorus* (Pyle 1997, Wolfe and Pyle 2011, Jenni and Winkler 2020), resulting in juvenile wing feathers being replaced by first alternate feathers. In these species definitive prealternate moults are also extensive, first prealternate moults often proceed in the same sequence as the preformative moults, and a sequence-based definition for each moult is maintained, despite variability in the sequence of some (but not all) prealternate moults, as mentioned above. Many other species reported to have more extensive prebreeding than postjuvenile moults (Jenni and Winkler 2020) would be interpeted under a sequence-based system as having suspended preformative moults.

A sequence-based definition can also be applied to non-passerine species that moult twice away from breeding grounds, such as those among divers, skuas, terns, and waders. Many species in these and other non-passerine families have well-documented overlapping prebasic and prealternate moults or, for example among terns and waders, overlapping prealternate and presupplmental moults (Pyle 2008, 2019), in each case feather-replacement sequence defines the initiation of each moult. It is thus not surprising that passerines can also show overlapping moults. Large non-passerine species that exhibit Staffealmauser or stepwise moults are generally considered to have incomplete prebasic moulting episodes as opposed to overlapping moults (Pyle 2005, 2008), although in some species (e.g., among Sulidae and Cathartidae) with rather continuous flight-feather replacement during prebreeding years, overlapping prebasic moults may also be inferred using a sequence-based approach (Pyle 2008, Chandler et al. 2010).

## CONCLUSIONS

I can think of no examples in which a sequence-based definition cannot be effectively applied to moult in migratory bird species. Although more study is needed on actual feather replacement sequence vs. resulting phonologies following partial moults (especially among wing coverts), I believe a sequence-based approach will eventually enable a clearer interpretation of moult strategies, especially when using an evolutionary approach to the definition of moults. Additionally, although some minor exceptions or perplexing situations might be predicted, solutions can be identified if definitions are based on the evolutionary histories of moult strategies. Due to the different bases for definition, moult terms under the life-history and H-P terminologies should not be considered synonyms of each other; however, should life-history definitions of postjuvenile, postbreeding, and prebreeding moults adopt a similar sequence-based approach to that described using H-P terminology here, moults away from breeding areas could be substantially clarified. In any case, those examining adaptive evolutionary factors that affect moult strategies should carefully define preformative/postjuvenile, prebasic/postbreeding, and prealternate/prebreeding moults. For such analyses I recommend using the sequence-based approach proposed here (cf. Guallar et al. 2021), while separate analyses considering feather replacement both pre- and post-migration (cf. Hall and Tullberg 2004, Delhey et al. 2020) may shed further light not only on environmental and life-history factors affecting the extents of moults overall, but on the those affecting variation in extents prior to suspension, at both individual and species levels.

